# Linking modulation of bio-molecular phase behaviour with collective interactions

**DOI:** 10.1101/2023.11.02.565376

**Authors:** Daoyuan Qian, Hannes Ausserwoger, William E. Arter, Rob M. Scrutton, Timothy J. Welsh, Tadas Kartanas, Niklas Ermann, Seema Qamar, Charlotte Fischer, Tomas Sneideris, Peter St George-Hyslop, Rohit V. Pappu, Tuomas P. J. Knowles

**Affiliations:** Centre for Misfolding Diseases, Yusuf Hamied Department of Chemistry, University of Cambridge, Lensfield Road, Cambridge, CB2 1EW, UK; Transition Bio Limited, Eastbrook, Shaftesbury Road, Cambridge, CB2 8DU, UK; Department of Medicine (Division of Neurology), Temerty Faculty of Medicine, University Health Network, University of Toronto, Toronto M5T 0S8, Canada; Department of Neurology, Columbia University, 710 West 168th Street, New York, NY 10032, USA; Department of Biomedical Engineering and Center for Biomolecular Condensates, Washington University in St. Louis, St. Louis, MO 63130, USA; Cavendish Laboratory, Department of Physics, University of Cambridge, J J Thomson Ave, Cambridge, CB3 0HE, UK

## Abstract

Bio-molecular condensates formed in the cytoplasm of cells are increasingly recognised as key spatiotemporal organisers of living matter, and are implicated in a wide range of functional or pathological processes. This discovery opens up a new avenue for condensate-based applications and a crucial step in controlling this process is to understand the underlying interactions driving condensate formation or dissolution. However, these condensates are highly multi-component assemblies and many inter-component interactions are present, rendering it difficult to identify key promoters of phase separation. In this work, we extend the recently formulated dominance analysis to modulations of condensate formation. By carrying out dilute phase concentration measurements of a single target solute, the theoretical framework allows one to deduce whether the modulator acts on the target solute or another unspecified, auxiliary solute, as well as the attractive/repulsive nature of the added interaction. This serve as a general guide towards deducing possible modulation mechanisms on the molecular level, which can be complemented by orthogonal measurements. As a case study, we investigate the modulation of G3BP1/RNA condensates by the small molecule suramin, and the dominance measurements point towards a dissolution mechanism where suramin acts on G3BP1 to disrupt G3BP1/RNA interactions, as confirmed by a diffusional sizing assay. Our approach and the dominance framework have a high degree of adaptability and can be applied in many other condensate-forming systems.

---

Since its first characterisation in 2009 [1], bio-molecular phase separation has emerged as an important process in cellular physiology. Many examples have been found where the formation of these liquid-like, protein- and/or RNA-rich droplets serves physiological functions in cells [2–5] or is implicated in neurodegenerative diseases [6, 7]. Furthermore, the liquid nature of the droplets leads to dynamic processes such as ripening [1, 8–11], wetting [12, 13] and ageing [14–17], which can participate in or interfere with normal biological processes [7, 18, 19]. There is thus a huge interest in understanding the driving forces of bio-molecular condensate formation and mechanisms of action of modulators. Efforts in the past has focused on either all-or-none dissolution effects [20, 21] or simply the saturation concentration for phase separation [22–24], and the main difficulty in establishing a generic framework for modulation studies lies in the multi-component nature of typical condensate systems. The presence of many solute species, including salt, pH, and possibly RNA and crowding molecules, means the underlying phase space is high-dimensional and theoretical implications of this complexity have remained poorly explored - until the recent formulation of the dominance framework [25].

In a general multi-component system, the large number of solute species leads to a complex network of interactions but the dominance formulation distils these microscopic details into intuitive experimental observables. The thermodynamic driving force for phase separation is the free energy difference Δf between the homogeneous, mixed state and the phase separated, de-mixed state. This phase separation leads to differences in concentrations across phases for all solutes, characterised by a tie line vector. Δf can be written as a sum of contributions by individual solute species [25] and the dominance *D*^*α*^ of a solute *α* quantifies its contribution relative to the whole system (Figure 1a). To apply the framework, we only require measurement of the dilute phase concentration of one solute, hereafter termed the ‘target solute’. Typically this will be the protein undergoing phase separation; we refer to the rest of solutes, including salt, RNA, crowders as ‘auxiliary’ (Figure 1b). We use simply *D* to denote the dominance value for the target solute unless stated otherwise. If the target solute constitutes an effective

**FIG. 1.**
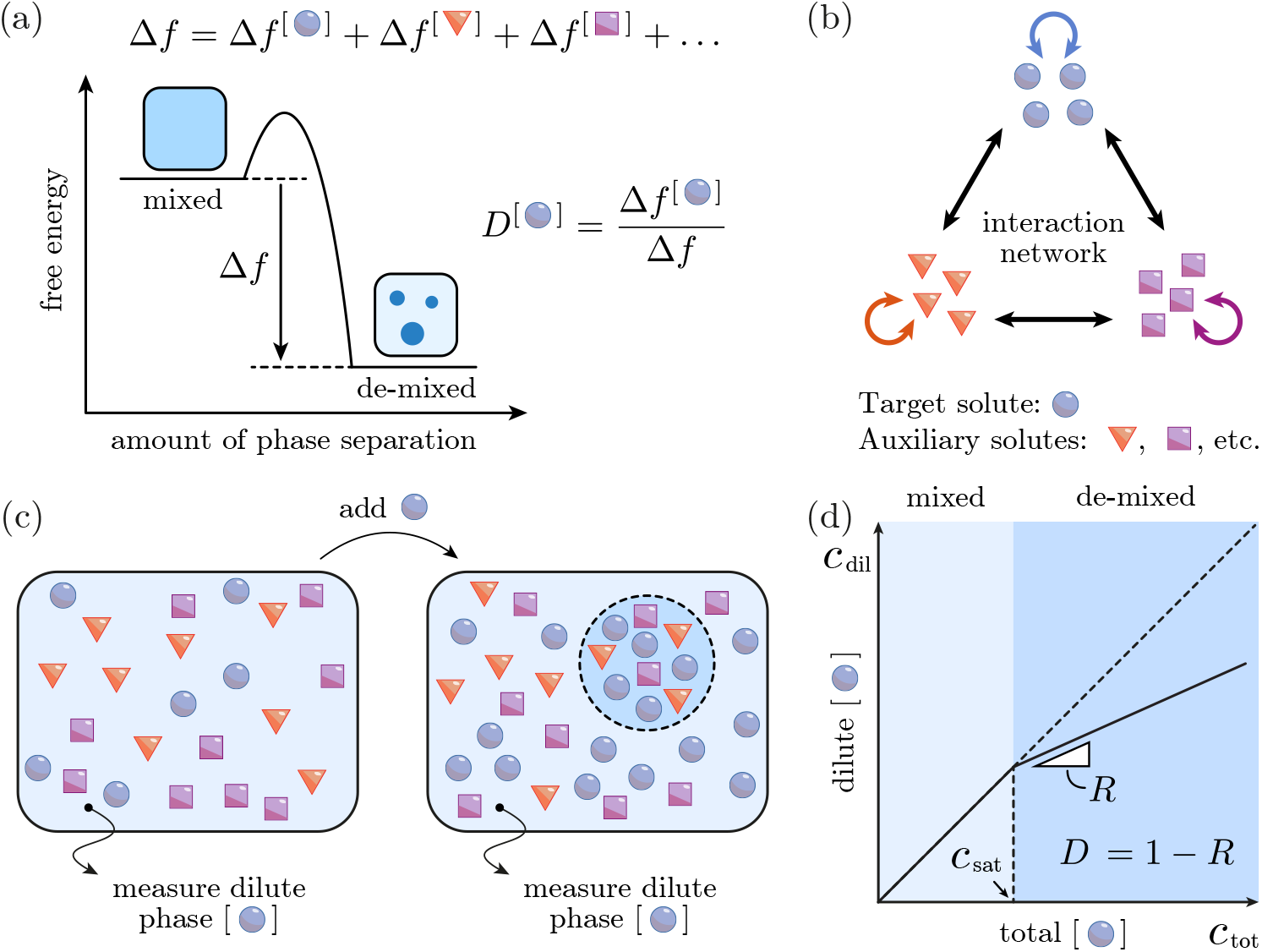
The dominance-based approach to study phase separation. (a) The phase separation process lowers the total free energy of the system by forming a dilute phase and a dense phase with an associated tie line vector. The free energy change Δ*f* can be written as a sum of contributions from each solute, and the dominance of a solute is its contribution relative to the total Δ*f* . (b) Phase separation occurs because of interactions among all solute species, resulting in a high-dimensional network. Among these solutes, we choose a target solute (blue), for which the dilute phase concentration is accessible. Other solutes are referred to as ‘auxiliary’ (red and purple). (c) Experimentally, a series of samples are prepared with varying concentrations of the target solute (*c*_tot_) while all other conditions are kept the same. For each sample the dilute phase concentration of the target solute (*c*_dil_) is measured. (d) In a *c*_tot_ − *c*_dil_ plot, the gradient *R* = 1 in the mixed region with no phase separation, and for *c*_tot_ *> c*_sat_ the sample enters the de-mixed region with *R <* 1 in general. The target dominance can be calculated as *D* = 1 − *R*. Diagonal dashed line indicates the trivial response with *R* = 1.

single-component system we will have *D* = 1, while multi-component characters can be detected if *D* lies between 0 and 1, and some of the auxiliary solutes can be contributing too. Experimental quantification of *D* is straight forward: it can be determined by measuring the dilute phase concentration of the target solute (*c*_dil_) in the phase-separated region, while changing the total concentration (*c*_tot_) of the target solute (Figure 1c). This is termed the ‘homotypic line-scan’ in [25] and is the focus of the current work. Its counterpart, ‘heterotypic line-scan’, can in principle be performed to gain the same insights [25, 26]. In a *c*_dil_ − *c*_tot_ plot, two regions can be identified: a mixed region with gradient 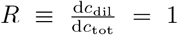 and a de-mixed region where *R <* 1 in general. The critical concentration *c*_sat_ marks the transition from the trivial response to the phase separation response. The dominance *D* of the target solute is *D* = 1 − *R*, evaluated right after *c*_sat_ (Figure 1d) [25]. The simple approach allows many existing datasets to be analysed and modulations of phase-separating systems can be studied too. Different types of modulations include mutations of a protein or additions of auxiliary solute molecules. The dominance and *c*_sat_ values can then be compared before and after the modulation effect is introduced, and experimentally this has already been done both *in vitro* and *in cellulo* (Table I). Comparative studies are performed for the poly(A) system under low or high salt conditions [25] (Table I 1, 2) and for the FUS system modulated by hexanediol [26] (Table I 3, 4), uncovering effects of electrostatic screening and disruption of hydrophobic interactions. In [27], an opto-genetic approach was used to express FUS, DDX4, and TAF15 protein constructs individually in cells, where protein-protein association is enhanced by exposing the cytoplasm to light. Homotypic line-scans are effectively performed with the proteins as target solutes, and mutations of these proteins are further introduced to decipher the sequence grammar of these protein systems. Here we analyse the data from [27]. For FUS, 5 tyrosine residues are replaced by 5 serine residues, reducing *c*_sat_ significantly while the *D* remains at a high value close to 1 (Table I 5, 6). This could be interpreted as removing the hydrophobic tyrosine reduces FUS-FUS attraction, although the free energy decrease is still mostly contributed by FUS alone. For DDX4, 4 phenylalanine residues are replaced by 4 alanine residues, also decreasing *c*_sat_ while the high noise in the *D* measurement only hints at a decrease in *D*, potentially due to increased involvement of other solutes, however more precise measurement is needed to confirm this (Table I 7, 8). For TAF15, by replacing 10 arginine with glutamine and 15 aspartic acid with serine residues, charge regulation of condensate formation is revealed as the less charged variant phase separates more easily at a lower *c*_sat_, accompanied by a likely increase in *D* (Table I 9, 10). Interpretations of above-mentioned results are intuitive since the modulations studied are relatively well-understood, but what if a small molecule is introduced to the system, interactions of which are unknown *a priori* ? To aid systematic analysis of dominance measurement data, here we further incorporate effects of modulations into the theoretical framework and derive rules governing changes on the observables *c*_sat_ and *D*, and then apply the results to a physiologically relevant G3BP1 system modulated by a small molecule suramin.

**TABLE I.**
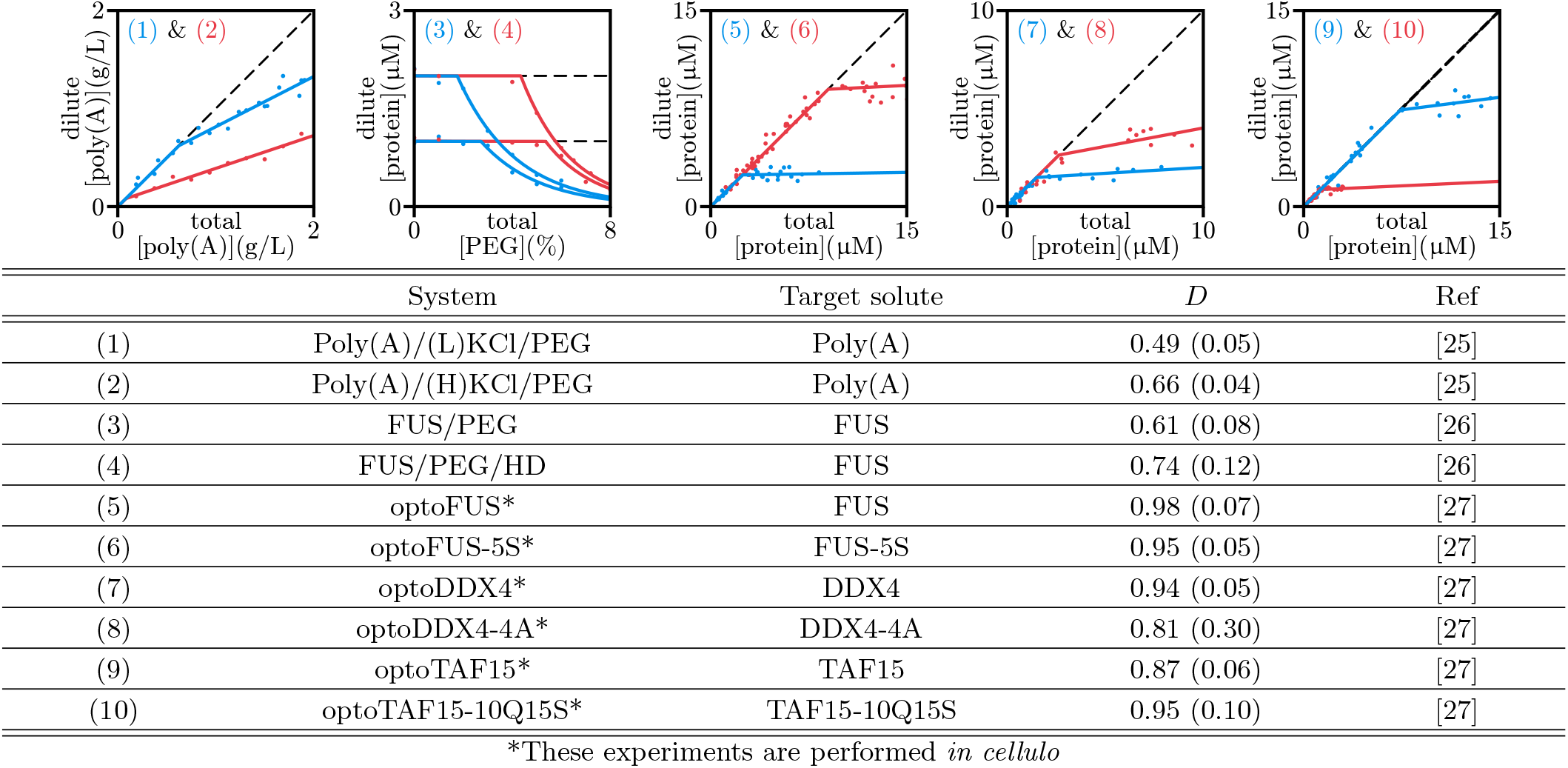
Dominance values for various phase separation systems. Top panels show the raw data (scatter) and the fitted response functions (solid lines) with trivial responses in black dashed lines. Systems (1) - (8) are analysed using the homotypic response and (9) - (10) are analysed using the heterotypic response. In (7) and (8), (L) and (H) stand for Low and High KCl concentrations respectively.

Denoting the initial free energy density as *f*, two generic types of modulations exist: changing *f* directly, for instance by changing the temperature, or shifting the homotypic line-scan to a different overall concentration composition, as in the case of adding modulators. It is however possible to show the equivalency between these two types of modulation (SI section I): we can incorporate the changing composition as a change in the ‘effective free energy’, rendering the theoretical treatment of modulations much simpler. The modulations can in principle be a complicated interaction network, while a simplified picture arises if we model the modulation as a sum of binary interactions. In doing so we can focus on the binary interaction with the largest magnitude, equivalent to focusing on the eigenvector of the perturbing interaction matrix with the largest absolute eigenvalue (SI section II). By performing homotypic line-scan with the target solute and studying how *c*_sat_ and *D* change under the modulation, insights into the perturbation can be gained. The change in *c*_sat_, denoted by Δ*c*_sat_, is easy to interpret. For an attractive perturbation, the modulation represents an enhancement of phase separation and we simply have Δ*c*_sat_ *<* 0. On the other hand, a repulsive perturbation suppresses condensate formation so Δ*c*_sat_ *>* 0 (SI section III). The change in *D* depends on both the attractive/repulsive nature of the perturbation interaction and how much it involves the target solute, compared to the tie line vector of the unperturbed system. If the perturbing interaction has a larger target solute content than the tie line vector we term it ‘target-aligned’, and otherwise we term it ‘auxiliary-aligned’. In the limit of the interaction involving only one solute, target-aligned means the perturbation is a homotypic interaction between the target solutes, while an auxiliary-aligned perturbation will be a homotypic interaction between solutes that are not the target solute. Theoretical analysis indicates that target-aligned enhancement makes the target dominance *D* larger and a target-aligned suppression makes *D* smaller, and similar observations can be made for auxiliary-aligned enhancement and suppression (SI section IV). Each modulation thus belongs to one of 4 broad classes: target enhancement/suppression, where attractive interactions involving the target solute gets stronger/weaker, and auxiliary enhancement/suppression, where attractive interactions involving one or more auxiliary solutes get stronger/weaker, respectively (Figure 2). This is the central result of the paper: by doing a simple dominance measurement at two different solvent conditions, the framework allows us to deduce which solute is most affected, and whether interactions involving that solute is getting more attractive or more repulsive.

**FIG. 2.**
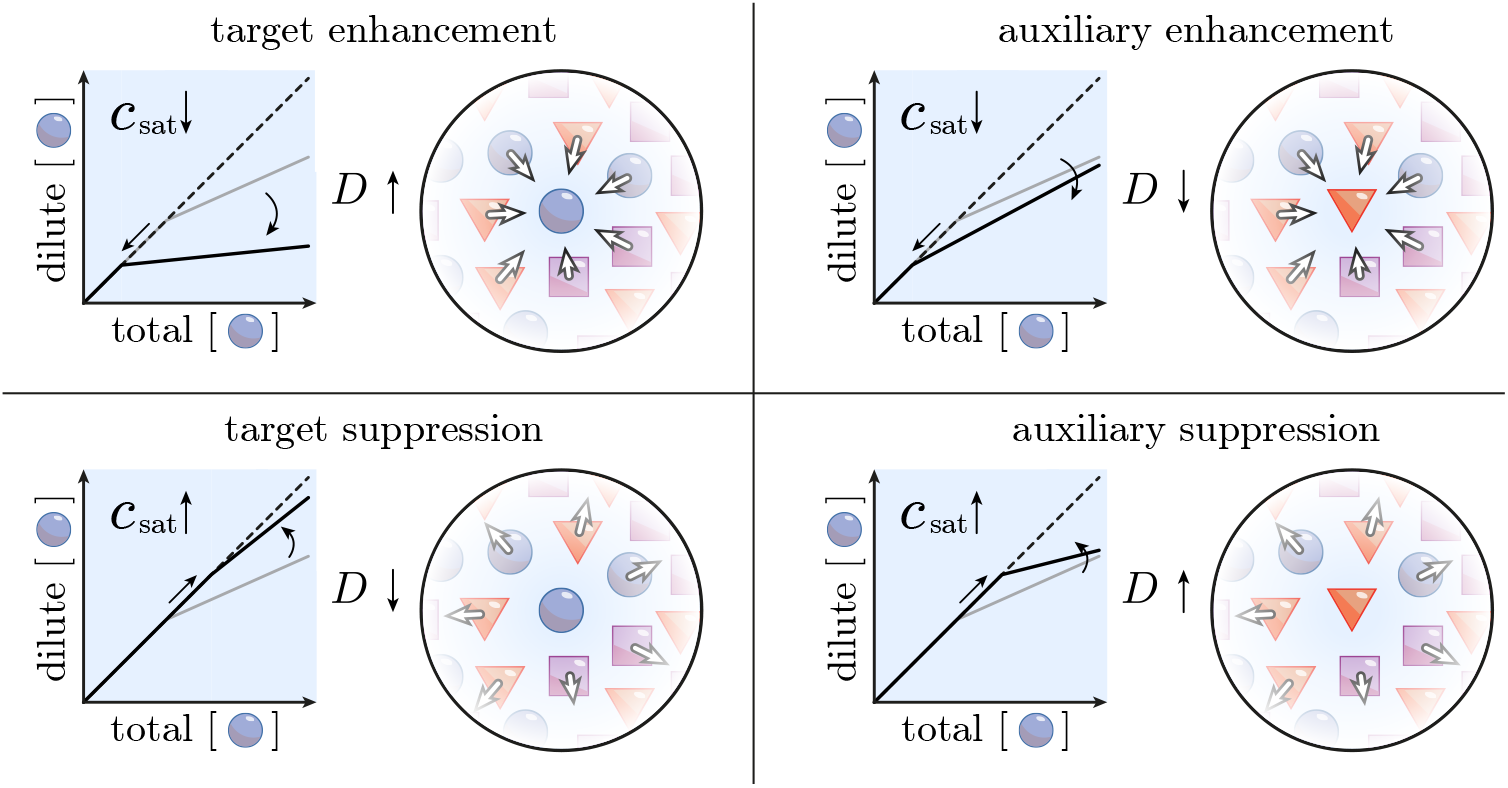
4 classes of mechanisms of modulation. We categorise each modulation according to whether it is an enhancement or suppression, and whether it is target-aligned or auxiliary-aligned. Each case is accompanied by a characteristic dilute phase target response function change (left panels, grey to black solid line). A simple interpretation of these modes is that the modulation brings about attractive interactions or repulsive interactions pertaining to one of the solute species (right panels, arrows going towards the central molecule indicates attraction and vice versa).

As a case study for the proposed modulation framework, we investigate the mechanism of action of the small molecule suramin on condensates formed by the protein G3BP1. G3BP1 is an RNA-binding protein central to stress granule formation [3, 4, 28] and is also involved in cancer progression [29, 30]. We use the PhaseScan platform [20] to sample the phase space along three axes: G3BP1, RNA and suramin, at a constant amount of PEG. For each sample point we quantify the total G3BP1, RNA, and suramin concentrations, as well as the dilute phase G3BP1 concentration and the presence/absence of condensates. G3BP1 is thus the target solute in the study. Experimental details can be found in SI section V. The phase boundaries from 3-dimensional PhaseScan showed dissolution action of both suramin and RNA, as is evident from taking 2-dimensional slices of the full dataset (Figure 3a, b). To perform dominance analysis, we section the data points into narrow windows of RNA and suramin concentrations and plot the dilute phase [G3BP1] against its total concentration (Figure 3b, inset), (SI section VI). We analyse *D* for a range of suramin and RNA values and *D* shows a decreasing trend when suramin is added at all RNA concentrations (Figure 3c). Combined with the dissolution action of suramin, we conclude suramin modulation is a target suppression so it acts on G3BP1 and weakens its interactions.

**FIG. 3.**
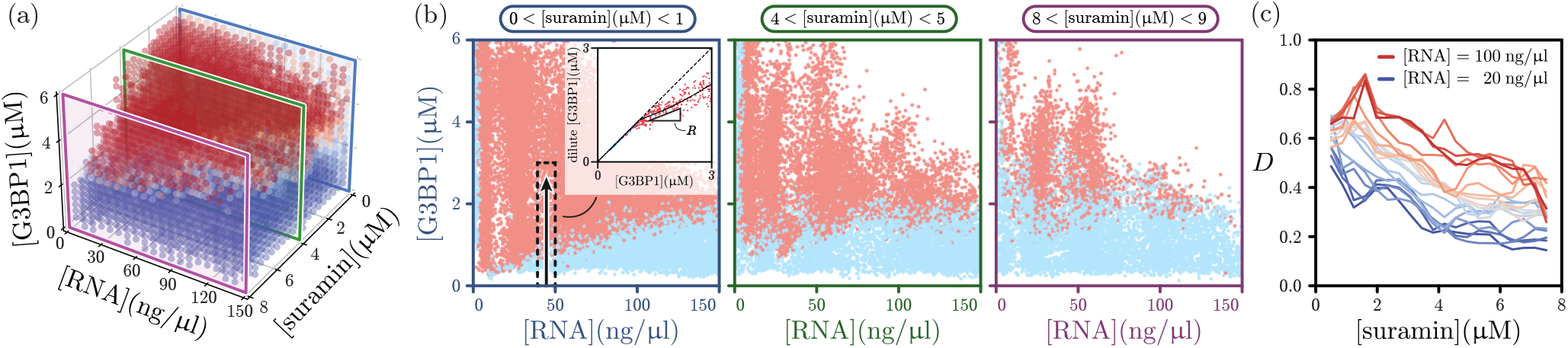
Dominance analysis applied to modulation of suramin on G3BP1/RNA condensates. (a) A 3-dimensional PhaseScan experiment is performed with varying total [G3BP1], [RNA] and [suramin]. Dilute phase [G3BP1] is measured in each droplet sample. Scatter points in the 3-dimensional plot are coarse-grained from the data, with red and blue corresponding to high and low phase separation probabilities respectively. To better visualise the data, we take 2-dimensional cross-sections along the suramin dimension (coloured rectangles). (b) Plots of constant [suramin] cross-sections. Each scatter point is a micro-droplet sample, with red and blue indicating presence and absence of condensates respectively. Left to right: sections of increasing total [suramin]. To calculate the dominance of G3BP1, we take a 1-dimensional data slice with a small window of [RNA] and [suramin] to produce the homotypic response curve (inset). (c) We perform the *D* evaluation at various [suramin] and [RNA] by taking multiple 1-D slices. Plotting *D* as a function of [suramin] shows a decreasing trend, so the suramin modulation is a target suppression. However, as [RNA] is increased, *D* increases as the system transitions towards a protein-limited situation, so addition of RNA is an auxiliary suppression. A plausible explanation of the latter is that at increasing levels of RNA, repulsion between RNA backbone charges gets stronger and thus RNA/RNA interactions weaken.

Apart from uncovering the mechanism of modulation of suramin, the dilute phase G3BP1 information can be used to deduce suramin’s effect on condensate composition too. In the low-RNA region of the G3BP1/RNA PhaseScan slice, the phase boundary is almost perpendicular to the RNA axis and we can use the dilute phase information to further deduce the tie line component ratio between G3BP1 and RNA. We assign alternating dark and light shades of red and blue to data points, and a different shade is used when dilute phase [G3BP1] crosses 1μM, 2μM, etc. This produces dilute phase band boundaries and in the mixed region, they are trivially flat but in the de-mixed region they have an upward slope at the low-RNA phase boundary (Figure 4a). The line-scan in this region shows *R*≈ 1 (Figure 4a, inset) and we use the geometrical formula for *D* to obtain *D* = 0.008 *±* 0.002, indicating a possibly large RNA dominance (SI section VII). The band boundaries arise from a dimensional reduction of tie lines onto the 2-D PhaseScan plane and can appear curved even if the tie lines are straight (SI section VII) [25] but near the low-RNA branch of the phase boundary, the gradient of these reduced tie lines, *K*, provides a good estimate for the G3BP1/RNA stoichiometry in the condensate (SI section VII). By focusing on the region where the band boundaries are parallel to one another (Figure 4a, dashed rectangle) we extract *K* as a function of suramin concentration. This shows a decrease in G3BP1 content relative to RNA, as suramin is added (Figure 4b).

**FIG. 4.**
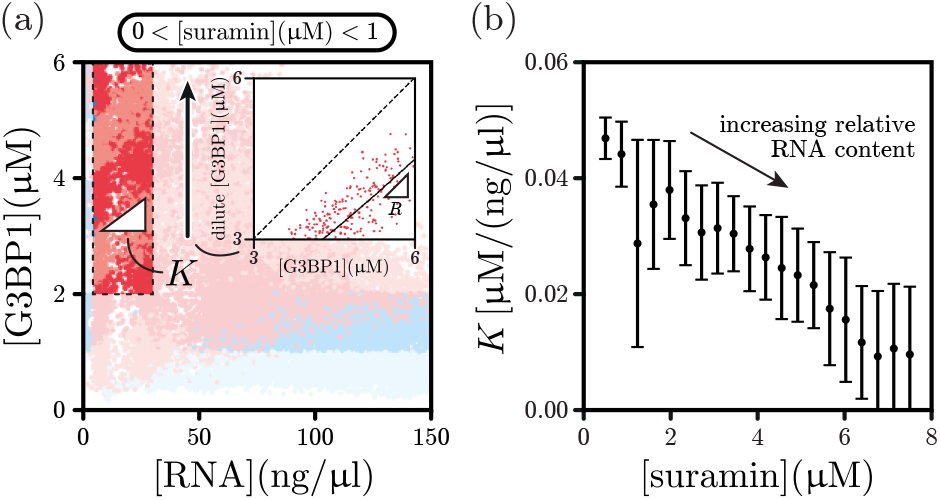
Tie line gradient estimation. (a) In the low-RNA regime (dashed rectangle region), RNA is likely the dominant solute and the gradient *K* of dilute phase band boundaries provides a good estimate for the G3BP1/RNA stoichiometry in condensates. Alternating shades of red and blue are produced by grouping data points based on its dilute phase G3BP1 concentrations. Inset: the line-scan in this regime shows a low *D* for G3BP1. (b) The stoichiometry decreases as suramin is added, indicating a increase in the amount of RNA relative to G3BP1 in condensates.

To deduce the exact mechanism of action of suramin on G3BP1 condensates we suppose the initial system has three main constituent solutes: G3BP1, RNA and PEG, with 3 homotypic and 3 heterotypic interactions. Upon addition of suramin, the suramin molecules can act on any of the 6 interactions. Since we have characterised the action as a target suppression, it has to weaken one of the 3 interactions that involve G3BP1. This however does not give a definitive answer on the interaction being modulated, so we perform a diffusional sizing assay [31, 32] to further elucidate the suramin action. We measure the hydrodyanmic radius *R*_h_ of G3BP1 at increasing RNA concentrations in the absence or presence of suramin. No PEG is added in the experiments so as to avoid phase separation. Experimental details can be found in SI section VIII. The control experiment without suramin (Figure 5a, red) showed an increase in G3BP1 *R*_h_ with increasing [RNA], reminiscent of nano-cluster formation observed in other condensate-forming systems [33, 34]. With the addition of 10μM of suramin (Figure 5a, blue), this behaviour disappears and G3BP1 *R*_h_ stays constant throughout the [RNA] range probed. Suramin thus suppresses G3BP1/RNA cluster formation, and we conclude it is the G3BP1/RNA interaction being modulated by suramin (Figure 5b).

**FIG. 5.**
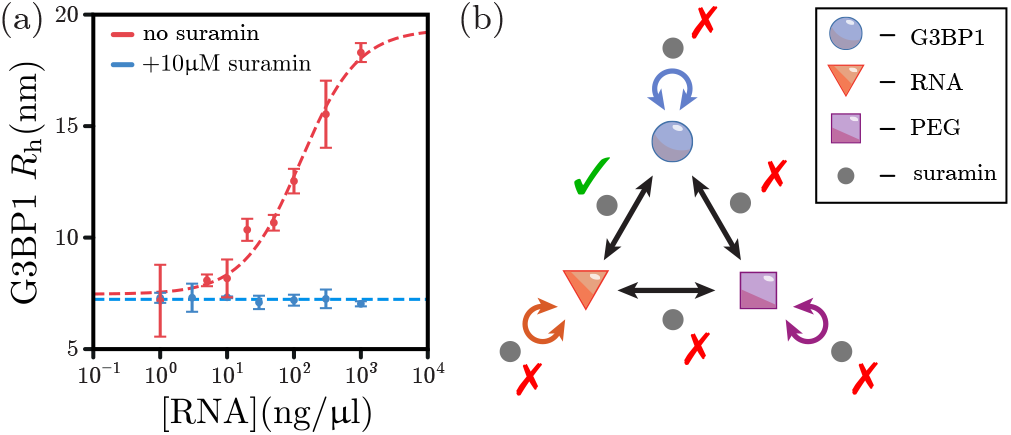
Sizing assay confirms mechanism of action. (a) Hydrodynamic radius *R*_h_ of G3BP1 as a function of [RNA]. Without suramin (red), the hydrodynamic radius of G3BP1 increases as RNA is added, indicating G3BP1/RNA binding. Upon addition of suramin, this behaviour is completely eliminated (blue). Red dashed line is a fitted binding curve. (b) Considering G3BP1, RNA and PEG as main phase separating solutes, suramin can possibly act on any of the 6 interactions (double arrows). Suramin having a target suppression effect means it acts on a G3BP1-related interaction, and performing the sizing assay allows us to further conclude that it is the G3BP1/RNA interaction being modulated.

The theoretical and experimental stories convey a central message: the dilute phase concentrations of solutes in phase separated systems contain a large amount of mechanistic information that has been overlooked so far, and under the dominance framework we have not only defined and measured multi-component characters of a phase separating system but also gained enough physical insight to deduce the mode of action of a modulator. The real merit in the framework is that it is completely general, oblivious of small-scale details of the system such as the molecular structure, charge, or even the size of the solute. The 4 classes of modulations serve as a generic guide towards unveiling mechanisms of action of modulators, together with their characteristic dominance measurement readouts. By treating all solutes on an equal footing this framework can be applied to any other phase separation system,as long as the dilute phase concentration of one solute can be determined post phase separation. We expect this approach to become central to modulation studies of bio-molecular condensates in the future.

## Acknowledgements

This study is supported by the Global Research Technologies Novo Nordisk A/S (H.A., T.P.J.K.), Harding Distinguished Postgraduate Scholar Programme (T.J.W.), Canadian Institutes of Health Research (Foundation Grant and Canadian Consortium on Neurodegeneration in Aging Grant to P.St.G.H.), US Alzheimer Society Zenith Grant ZEN-18-529769 (P.St.G.H.), Wellcome Trust Collaborative Award 203249/Z/16/Z (P.St.G.H., T.P.J.K.), and the European Research Council under the European Union’s Seventh Horizon 2020 research and innovation program through the ERC grant DiProPhys, agreement ID 101001615 (T.P.J.K.).

