## Supplementary Information for "Linking modulation of bio-molecular phase behaviour with collective interactions"

### Supplementary Information for: Deciphering modulations of bio-molecular phase separation

**Notations.** In this supplementary information document we follow conventions from differential geometry, where summation over repeated greek indices from 1 to $N$ is assumed unless otherwise stated. In cases where a summation is not implied, or if an index correspond to a scalar quantity instead of a vector component we simply put a round bracket around the index, e.g. $(\alpha)$. We keep the discussion general by considering all solute species $\alpha = 1, 2, \ldots, N$, denoting their total concentrations as $\phi^\alpha$, and dilute and dense phase concentrations as $\psi_-^\alpha$ and $\psi_+^\alpha$ respectively. The saturation concentration $c_{\text{sat}}$ in the context of homotypic line-scan is denoted $\phi_c^{(\alpha)}$ for a single index $(\alpha)$, and represents the $\psi_-^{(\alpha)}$ value given a set of $\psi_-^{(\beta)}$ for $(\beta) \neq (\alpha)$.

#### I. EQUIVALENCY BETWEEN TWO MODULATION TYPES

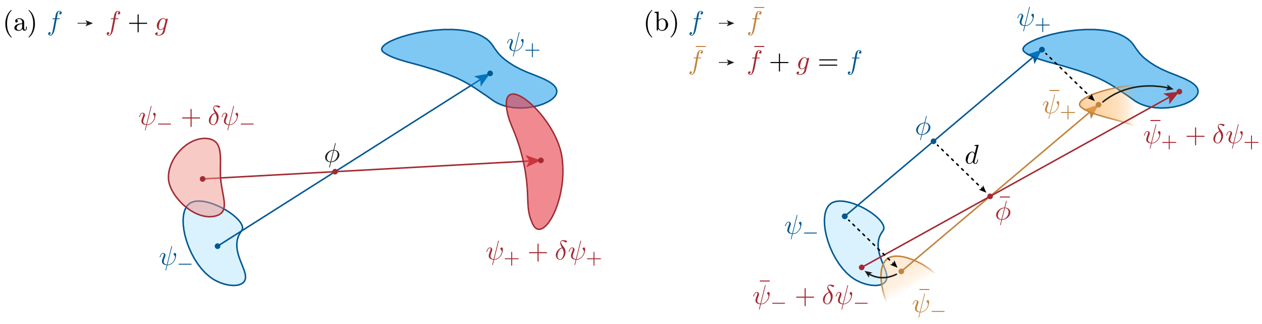


FIG. S1. **Free energy change as a fundamental mechanism of modulation.** (a) Starting with a free energy density $f$, the point $\phi^\alpha$ phase-separates into dilute and dense phase points $\psi_-^\alpha$ and $\psi_+^\alpha$, connected by a tie line (blue). Under a free energy change $f \to f + g$ with $\phi^\alpha$ staying the same, dilute and dense phase compositions move by $\delta\psi_-^\alpha$ and $\delta\psi_+^\alpha$ respectively (red). (b) A change in total composition $\phi^\alpha \to \bar{\phi}^\alpha = \phi^\alpha + d^\alpha$ can be treated as a free energy change. Translating $\phi^\alpha$ to $\bar{\phi}^\alpha$ is equivalent to a coordinate transformation by $d^\alpha$ with a free energy change $f \to \bar{f}$ such that the dilute and dense phases stay the same relative to each other (blue to yellow). To recover the real effect of the shift in $\phi^\alpha$ we apply an additional free energy change $\bar{f} \to \bar{f} + g = f$ such that the original free energy is recovered (yellow to red).

Assuming a system is associated with a free energy density $f = f(\phi)$ and an average total composition $\phi^\alpha$, we study modifications of $f$ such that $f \to f + g$ where $g$ represents a perturbation to $f$. Denoting the initial equilibrium dilute and dense phase compositions as $\psi_-^\alpha$ and $\psi_+^\alpha$ respectively, the perturbation results in changes in equilibrium compositions by $\delta\psi_-^\alpha$ and $\delta\psi_+^\alpha$ (Figure S1a). This can be the model for a temperature change. Modulation by addition of solute species, on the other hand, is not directly modifying $f$ but rather shifting $\phi^\alpha$ in the phase space. We can however model this still as a free energy change by treating the change in $\phi^\alpha$ as a coordinate transformation. Denoting the initial equilibrium points as $\phi^\alpha$, $\psi_-^\alpha$, and $\psi_+^\alpha$ as before, and the new total composition as $\bar{\phi}^\alpha$, the difference

$$d^\alpha \equiv \bar{\phi}^\alpha - \phi^\alpha \tag{S.1}$$

is a constant vector along which all points are shifted. Suppose that associated with this new coordinate there is a free energy $\bar{f}$ such that $\bar{f}(\bar{\phi}) = f(\phi)$. Under $\bar{f}$, the dilute and dense phase equilibrium compositions are simply shifted by $d^\alpha$: $\bar{\psi}_\pm = \psi_\pm + d^\alpha$. Next, to recover the physics of changing $\phi^\alpha$ we have to compute the equilibrium with $f$ instead of $\bar{f}$. This can be modelled as changing $\bar{f}$ by a function $g$: $\bar{f} \to \bar{f} + g$ such that $\bar{f}(\bar{\phi}) + g(\bar{\phi}) = f(\bar{\phi})$. Recall that $\bar{f}(\bar{\phi}) = f(\phi)$ and $\bar{\phi}^\alpha = \phi^\alpha + d^\alpha$ we arrive at an expression for $g$:

$$g(\bar{\phi}) = f(\bar{\phi}) - f(\bar{\phi} - d). \tag{S.2}$$

Under this change, suppose the equilibrium points $\bar{\psi}_\pm^\alpha$ are shifted by $\delta\psi_\pm^\alpha$, so the total response in $\psi_\pm^\alpha$ is $d^\alpha + \delta\psi_\pm^\alpha$ (Figure S1b). As such, both classes of modulations can be treated as a free energy perturbation and in derivations below we do not make this distinction unless otherwise explained.

#### II. UNIVERSAL TWO-BODY INTERACTION REPRESENTED BY A SINGULAR STOICHIOMETRY

Two-body interactions can be modelled generally as a quadratic perturbation function

$$g = \frac{1}{2}\Lambda_{\alpha\beta}\phi^{\alpha}\phi^{\beta} \tag{S.3}$$

with a symmetric interaction matrix $\Lambda_{\alpha\beta}$. This matrix can be diagonalised into $N$ eigenstates, generically written as

$$\Lambda_{\alpha\beta} = \sum_{(i)=1}^{N} \lambda^{(i)}\eta_{\alpha}^{(i)}\eta_{\beta}^{(i)} \tag{S.4}$$

and thus we can decompose the overall perturbation as a series of singular perturbations of the form

$$g = \frac{1}{2}\lambda\eta_{\alpha}\eta_{\beta}\phi^{\alpha}\phi^{\beta} \tag{S.5}$$

where now $\lambda$ determines the strength and nature of the interaction and $\eta_{\alpha}$ represents the relative solute content for this mode. For an attractive interaction we have $\lambda < 0$, and $\lambda > 0$ for a repulsive interaction. $\eta_{\alpha}$ is the singular stoichiometry in the main text. In cases where $\eta_{\alpha}$ is completely aligned with a solute axis, this then represents a homotypic interaction.

#### III. DEDUCING THE NATURE OF THE PERTURBATION FROM $\phi_{\mathrm{c}}$ MEASUREMENT

The system at equilibrium is defined by the set of equations arising from equilibration of chemical potentials, the osmotic pressure, and mass balance:

$$\begin{aligned} \partial_{\alpha}f(\psi_{-}) =& \partial_{\alpha}f(\psi_{+}), && \text{(S.6a)} \\ f(\psi_{-}) - \psi_{-}^{\alpha}\partial_{\alpha}f(\psi_{-}) =& f(\psi_{+}) - \psi_{+}^{\alpha}\partial_{\alpha}f(\psi_{+}), && \text{(S.6b)} \\ \phi^{\alpha} =& \psi_{-}^{\alpha} + vk^{\alpha}. && \text{(S.6c)} \end{aligned}$$

where the tie line vector is $k^{\alpha} \equiv \psi_{+}^{\alpha} - \psi_{-}^{\alpha}$. The set of perturbations is given by

$$\begin{aligned} f \rightarrow& f + g \\ \psi_{\pm}^{\alpha} \rightarrow& \psi_{\pm}^{\alpha} + \delta\psi_{\pm}^{\alpha} \\ v \rightarrow& v + \delta v \end{aligned} \tag{S.7}$$

and we assume the responses are of order $\mathcal{O}(\epsilon)$ and that $f$ and $g$ are locally at most quadratic. This means we ignore derivatives of the kind $\partial_{\alpha}\partial_{\beta}\partial_{\gamma}f$ or $\partial_{\alpha}\partial_{\beta}\partial_{\gamma}g$. For now we keep the form of $g$ general instead of using the singular form. Under the free energy change $g$, changes in equilibrium chemical potentials and osmotic pressure are, to first order,

$$\begin{aligned} \delta\left[\partial_{\alpha}f(\psi_{-})\right] =& + \delta\psi_{-}^{\beta}\partial_{\alpha}\partial_{\beta}f(\psi_{-}) + \partial_{\alpha}g(\psi_{-}) + \delta\psi_{-}^{\beta}\partial_{\alpha}\partial_{\beta}g(\psi_{-}) + \mathcal{O}(\epsilon^{2}) \\ \delta\left[f(\psi_{-}) - \psi_{-}^{\alpha}\partial_{\alpha}f(\psi_{-})\right] =& + \delta\psi_{-}^{\alpha}\partial_{\alpha}f(\psi_{-}) + g(\psi_{-}) + \delta\psi_{-}^{\alpha}\partial_{\alpha}g(\psi_{-}) \\ & - \left(\psi_{-}^{\alpha} + \delta\psi_{-}^{\alpha}\right)\left[\partial_{\alpha}f(\psi_{-}) + \delta\psi_{-}^{\beta}\partial_{\alpha}\partial_{\beta}f(\psi_{-}) + \partial_{\alpha}g(\psi_{-}) + \delta\psi_{-}^{\beta}\partial_{\alpha}\partial_{\beta}g(\psi_{-}) + \mathcal{O}(\epsilon^{2})\right] \\ & + \psi_{-}^{\alpha}\partial_{\alpha}f(\psi_{-}) \\ =& + g(\psi_{-}) \\ & - \psi_{-}^{\alpha}\left[\partial_{\alpha}f(\psi_{-}) + \delta\psi_{-}^{\beta}\partial_{\alpha}\partial_{\beta}f(\psi_{-}) + \partial_{\alpha}g(\psi_{-}) + \delta\psi_{-}^{\beta}\partial_{\alpha}\partial_{\beta}g(\psi_{-})\right] \\ & + \psi_{-}^{\alpha}\partial_{\alpha}f(\psi_{-}) + \mathcal{O}(\epsilon^{2}) \\ =& + g(\psi_{-}) - \psi_{-}^{\alpha}\delta[\partial_{\alpha}f(\psi_{-})] + \mathcal{O}(\epsilon^{2}) \end{aligned} \tag{S.8}$$

and it is easy to verify that constant or linear contributions of $g$ do not induce any response. Equilibrium of osmotic pressure requires

$$\begin{aligned} g(\psi_{+}) - g(\psi_{-}) =& k^{\alpha}\delta[\partial_{\alpha}f(\psi_{-})] \\ g(\psi_{+}) - g(\psi_{-}) - k^{\alpha}\partial_{\alpha}g(\psi_{-}) =& \delta\psi_{-}^{\alpha}k^{\beta}[\partial_{\alpha}\partial_{\beta}f(\psi_{-}) + \partial_{\alpha}\partial_{\beta}g(\psi_{-})] \end{aligned} \tag{S.9}$$

Using the mass balance Eq. (S.6c)

$$\delta\psi_-^\alpha = -\left(\delta v k^\alpha + v\delta k^\alpha\right) = -\delta(vk^\alpha), \tag{S.10}$$

we get

$$\begin{aligned} g(\psi_+) - g(\psi_-) - k^\alpha\partial_\alpha g(\psi_-) &= -\,\delta\left(vk^\alpha\right) k^\beta[\partial_\alpha\partial_\beta f(\psi_-) + \partial_\alpha\partial_\beta g(\psi_-)] \\ vg(\psi_+) - vg(\psi_-) - vk^\alpha\partial_\alpha g(\psi_-) &= -\,\delta\left(vk^\alpha\right)(vk^\beta)[\partial_\alpha\partial_\beta f(\psi_-) + \partial_\alpha\partial_\beta g(\psi_-)] \\ vg(\psi_+) - vg(\psi_-) - vk^\alpha\partial_\alpha g(\psi_-) - \frac{1}{2}vk^\alpha vk^\beta\partial_\alpha\partial_\beta g(\psi_-) &= -\,\delta\left(vk^\alpha\right)(vk^\beta)[\partial_\alpha\partial_\beta f(\psi_-) + \partial_\alpha\partial_\beta g(\psi_-)] - \frac{1}{2}vk^\alpha vk^\beta\partial_\alpha\partial_\beta g(\psi_-) \end{aligned} \tag{S.11}$$

now the right hand side is $\delta\Delta f$, recall the expression of $\Delta f$ [1]:

$$\Delta f = -\frac{1}{2}vk^\alpha vk^\beta\partial_\alpha\partial_\beta f(\psi_-), \tag{S.12}$$

the left hand can be re-arranged to be

$$\begin{aligned} \text{L. H. S.} =& vg(\psi_+) - g(\psi_-) + g(\psi_-) - vg(\psi_-) - vk^\alpha\partial_\alpha g(\psi_-) - \frac{1}{2}vk^\alpha vk^\beta\partial_\alpha\partial_\beta g(\psi_-) \\ =&(1-v)g(\psi_-) + vg(\psi_+) - g(\phi) \\ \equiv&\Delta g \end{aligned} \tag{S.13}$$

so we have $\Delta g = \delta\Delta f$. $\Delta g$ represents the extra stabilisation energy due to $g$ at the original compositions, and the response of the system fully captures this gain (or loss).

To work out $\delta v$, we expand out $\delta(vk)$ in the first line of Eq. (S.11),

$$g(\psi_+) - g(\psi_-) - k^\alpha\partial_\alpha g(\psi_-) = -\left(\delta v k^\alpha + v\delta k^\alpha\right) k^\beta[\partial_\alpha\partial_\beta f(\psi_-) + \partial_\alpha\partial_\beta g(\psi_-)] \tag{S.14}$$

and in the limit of $v = 0$ we have

$$\delta v = -\frac{g(\psi_+) - g(\psi_-) - k^\alpha\partial_\alpha g(\psi_-)}{k^\alpha[\partial_\alpha\partial_\beta f(\psi_-) + \partial_\alpha\partial_\beta g(\psi_-)]k^\beta} \tag{S.15}$$

where the numerator represents an energy difference between the dense phase and apparent dilute phase, equal to 0 if $\psi_\pm$ satisfies the common tangent construction with respect to $g$. The denominator consists of Hessians of $f$ and $g$, and since $\psi_\pm$ is locally stable all eigenvalues of $\partial_\alpha\partial_\beta f(\psi_-)$ are positive, leading to a positive-definite $k^\alpha k^\beta\partial_\alpha\partial_\beta f(\psi_-)$. If the perturbation $g$ is weak enough we thus expect the denominator to be positive in general. It is interesting to note that $\delta v$ does not depend on the Hessians evaluated at $\psi_+$, as the common tangent construction ensures the information contained there has been fully captured by the dilute phase.

Now we investigate the effect of a singular perturbation. Substituting in a singular $g$

$$g(\phi) \equiv \frac{1}{2}\lambda\eta_\alpha\eta_\beta\phi^\alpha\phi^\beta \tag{S.16}$$

gives

$$\delta v = -\frac{1}{2}\frac{\lambda(\eta_\alpha k^\alpha)^2}{(k^\alpha H_{\alpha\beta}k^\beta) + \lambda(\eta_\alpha k^\alpha)^2} \tag{S.17}$$

where $H_{\alpha\beta} \equiv \partial_\alpha\partial_\beta f(\psi_-)$. As such, the sign of $\delta v$ depends solely on the attractive/repulsive nature of $g$, given the perturbation is weaker than the existing Hessian. This expression is arrived at by taking the $v \to 0$ limit, so we get an expression for the dilute phase response

$$\delta\psi_-^\alpha = -k^\alpha\delta v = (\lambda k^\alpha)\left[\frac{1}{2}\frac{(\eta_\alpha k^\alpha)^2}{(k^\alpha H_{\alpha\beta}k^\beta) + \lambda(\eta_\alpha k^\alpha)^2}\right] \tag{S.18}$$

where the square bracket is always positive, so the sign of $\delta\psi_-^\alpha$ depends solely on that of $\lambda k^\alpha$. In experimental line-scan, however, it is the critical concentration $\phi_c^{(\alpha)}$ that is characterised for a target solute $(\alpha)$, and to relate $\delta\phi_c^{(\alpha)}$ to $\delta\psi_-^{(\alpha)}$ we use the relation

$$\delta\phi_c^{(\alpha)} = \frac{\delta\psi_-^{(\alpha)}}{D^{(\alpha)}} \tag{S.19}$$

which can be easily derived by observing the geometry of the homotypic response function. The sign of $\delta\phi_c^{(\alpha)}$ is thus given by

$$\mathrm{sgn}\left[\delta\phi_c^{(\alpha)}\right] = \mathrm{sgn}\left[\lambda\right]\mathrm{sgn}\left[k^{(\alpha)}\right]\mathrm{sgn}\left[D^{(\alpha)}\right] \tag{S.20}$$

so given a normal solute $(\alpha)$ with $0 < D^{(\alpha)} < 1$, that also enriches in condensates $[k^{(\alpha)} > 0]$, adding an attractive interaction $(\lambda < 0)$ leads to a decrease in $\phi_c^{(\alpha)}$ and vice versa. We can thus broadly categorise the modulation effects as either an enhancement $(\lambda < 0)$ or suppression $(\lambda > 0)$ of phase-separation, interpretable from $\phi_c^{(\alpha)}$. It is worth noting that this result does not depend on $\eta_\alpha$ at all - and next we explore how the alignment of $\eta_\alpha$ affects the dominance $D^{(\alpha)}$.

#### IV. DEDUCING THE SINGULAR STOICHIOMETRY ALIGNMENT FROM $D$ MEASUREMENT

Using the singular form of $g$

$$g(\phi) \equiv \frac{1}{2}\lambda\eta_\alpha\eta_\beta\phi^\alpha\phi^\beta, \tag{S.21}$$

equilibrium of chemical potentials written using Eqs. (S.8) become

$$\delta\psi_-^\beta\partial_\alpha\partial_\beta f(\psi_-) + \lambda\psi_-^\beta\eta_\alpha\eta_\beta + \delta\psi_-^\beta\lambda\eta_\alpha\eta_\beta = \delta\psi_+^\beta\partial_\alpha\partial_\beta f(\psi_+) + \lambda\psi_+^\beta\eta_\alpha\eta_\beta + \delta\psi_+^\beta\lambda\eta_\alpha\eta_\beta. \tag{S.22}$$

To tackle Eqs. (S.22) we write, as before,

$$H_{\alpha\beta} \equiv \partial_\alpha\partial_\beta f(\psi_-) = \partial_\alpha\partial_\beta f(\psi_+) - \delta H_{\alpha\beta} \tag{S.23}$$

with $\delta H_{\alpha\beta} \equiv \partial_\alpha\partial_\beta f(\psi_+) - \partial_\alpha\partial_\beta f(\psi_-)$. We have

$$\delta\psi_-^\beta H_{\alpha\beta} + \lambda\psi_-^\beta\eta_\alpha\eta_\beta + \delta\psi_-^\beta\lambda\eta_\alpha\eta_\beta = \delta\psi_+^\beta(H_{\alpha\beta} + \delta H_{\alpha\beta}) + \lambda\psi_+^\beta\eta_\alpha\eta_\beta + \delta\psi_+^\beta\lambda\eta_\alpha\eta_\beta, \tag{S.24}$$

so, using $\psi_+^\alpha = k^\alpha + \psi_-^\alpha$,

$$\begin{aligned} 0 &= \delta k^\beta H_{\alpha\beta} + \lambda k^\beta\eta_\alpha\eta_\beta + \delta k^\beta\lambda\eta_\alpha\eta_\beta + \delta\psi_+^\beta\delta H_{\alpha\beta} \\ &= \delta k^\beta(H_{\alpha\beta} + \lambda\eta_\alpha\eta_\beta + \delta H_{\alpha\beta}) + \lambda k^\beta\eta_\alpha\eta_\beta + \delta\psi_-^\beta\delta H_{\alpha\beta} \end{aligned} \tag{S.25}$$

Using the mass balance Eq. (S.6c) and taking the limit of $v \to 0$ we have

$$\delta\psi_-^\alpha = -\delta v k^\alpha, \tag{S.26}$$

and Eqs. (S.25) gives

$$\delta k^\beta(H_{\alpha\beta} + \delta H_{\alpha\beta} + \lambda\eta_\alpha\eta_\beta) = -k^\beta(\lambda\eta_\alpha\eta_\beta - \delta v\delta H_{\alpha\beta}) \tag{S.27}$$

and we need to compute the inverse of the matrix on the left hand side to solve for $\delta k^\alpha$. To 0-th order we let $\delta H_{\alpha\beta} = 0$, and inverting $(H_{\alpha\beta} + \lambda\eta_\alpha\eta_\beta)$, we have

$$\delta k^\alpha = -\lambda k^\beta\eta_\beta\eta_\gamma\left[\left(H^{-1}\right)^{\gamma\alpha} - \frac{\lambda\left(H^{-1}\right)^{\sigma\gamma}\eta_\sigma\left(H^{-1}\right)^{\zeta\alpha}\eta_\zeta}{1 + \lambda\left(H^{-1}\right)^{\sigma\zeta}\eta_\sigma\eta_\zeta}\right] = -\left[\frac{\lambda k^\beta\eta_\beta}{1 + \lambda\left(H^{-1}\right)^{\sigma\zeta}\eta_\sigma\eta_\zeta}\right]\eta_\gamma\left(H^{-1}\right)^{\gamma\alpha}. \tag{S.28}$$

where $\left(H^{-1}\right)^{\alpha\beta}$ is the inverse of $H_{\alpha\beta}$, representing the local instability. Eq. (S.28) contains information on how the tie line vector changes under the influence of $g$. To build a geometrical understanding of $\delta k^\alpha$ relative to all the other vectors, first note that $n_\alpha$ and $\eta_\alpha$ are defined up to an arbitrary multiplicative constant, and only $k^\alpha$ has a well-defined sign: it points from the dilute phase towards the dense phase. We choose the sign convention for $\eta_\alpha$ such that $\eta_\alpha k^\alpha > 0$. Furthermore, the denominator of Eq. (S.28) is positive for $\lambda > 0$, since all eigenvalues of $H_{\alpha\beta}$ and thus of $\left(H^{-1}\right)^{\alpha\beta}$ are positive. The denominator is negative only in the case of a very attractive $g$ at $\lambda < -1/\left[\left(H^{-1}\right)^{\alpha\beta}\eta_\alpha\eta_\beta\right]$, at which point the locally stable dense and dilute phase points become locally unstable. We assume the perturbation is either repulsive, or in the case of an attractive $g$, weak enough so the denominator of Eq. (S.28) is always positive. A physical interpretation of Eq. (S.28) is the following. The tie line vector $k^\alpha$ responds to $g$ primarily by aligning itself parallel or anti-parallel to $\eta_\alpha$ in cases of additional attraction ($\lambda < 0$) or repulsion ($\lambda > 0$) of solute species enriched in the dense phase (this is imposed by choosing $k^\alpha\eta_\alpha > 0$). The degree of alignment however is not exactly parallel to $\eta_\alpha$ but is modulated by the local stability of the equilibrium points, and this is effected by multiplying $\eta_\alpha$ by the local instability matrix $\left(H^{-1}\right)^{\alpha\beta}$. $\delta k^\alpha$ can thus be expressed as, up to a positive multiplicative constant,

$$\delta k^\alpha \propto -\lambda\eta_\gamma(H^{-1})^{\gamma\alpha} \tag{S.29}$$

and to simplify this further we can approximate the Hessian to be isotropic, with $H_{\alpha\beta} \propto \mathbb{1}_{\alpha\beta}$ and $(H^{-1})^{\alpha\beta} \propto \mathbb{1}^{\alpha\beta}$ up to positive multiplicative constants. This gives

$$\delta k^\alpha \propto -\lambda\eta^\alpha \tag{S.30}$$

so the perturbing singular stoichiometry is proportional to $\delta k^\alpha$ when the local Hessian is isotropic. Now we examine $\delta n_\alpha$ and $\delta D^{(\alpha)}$ under the perturbation. $n_\alpha$ is given by

$$n_\alpha \propto Ak^\beta H_{\alpha\beta} \tag{S.31}$$

for some constant $A$. Under the perturbation and ignoring third and higher-order derivatives of $f$, noting Eq. (S.25),

$$\begin{aligned}\delta n_\alpha =& A(\delta k^\beta H_{\alpha\beta} + \delta k^\beta\lambda\eta_\alpha\eta_\beta + k^\beta\lambda\eta_\alpha\eta_\beta)\\ =& -A(\delta\psi_-^\beta\delta H_{\alpha\beta} + \delta k^\beta\delta H_{\alpha\beta})\\ =& 0 \quad \text{by setting } \delta H \to 0.\end{aligned} \tag{S.32}$$

So $n_\alpha$ is constant. This is not true in general when higher-order derivatives of $f$ are significant, but we leave these more subtle scenarios to future work. The response of the target dominance $D^{(\alpha)} = \frac{n_{(\alpha)}k^{(\alpha)}}{n_\gamma k^\gamma}$ [1] is $\delta D^{(\alpha)} = D^{(\alpha)}\left[\frac{\delta k^{(\alpha)}}{k^{(\alpha)}} - \frac{n_\sigma\delta k^\sigma}{n_\gamma k^\gamma}\right]$. To interpret this, we observe that $n_\gamma$ and the axis $\delta_\gamma^{(\alpha)}$ define a 2-D plane, and only components of $k^\alpha$ and $\delta k^\alpha$ on this plane affects $\delta D^{(\alpha)}$. To put it in mathematical terms, we can rotate the coordinate system such that the $(\alpha)$ axis is left unchanged, and align another axis $(\beta)$ such that $n_\alpha$ only has non-zero components in the $(\alpha)$ and $(\beta)$ directions. This means for an arbitrary $x^\alpha$ we can write $n_\alpha x^\alpha = n_{(\alpha)}x^{(\alpha)} + n_{(\beta)}x^{(\beta)}$ so we have a simpler form for $\delta D^{(\alpha)}$:

$$\delta D^{(\alpha)} = D^{(\alpha)}\left[1 - D^{(\alpha)}\right]\left[\frac{\delta k^{(\alpha)}}{k^{(\alpha)}} - \frac{\delta k^{(\beta)}}{k^{(\beta)}}\right]. \tag{S.33}$$

The sign of $\delta D^{(\alpha)}$ is apparent after choosing the $(\alpha)$ and $(\beta)$ axes. For a typical solute $(\alpha)$ with $0 < D^{(\alpha)} < 1$ the sign of $\delta D^{(\alpha)}$ is entirely determined by whether $\delta k^\alpha$ is aligned closer to the $(\alpha)$ or $(\beta)$ axis in the $(\alpha) - (\beta)$ plane relative to the existing tie line vector $k^\alpha$ (Figure S2). Using the simple expression for $\delta k^\alpha$ in Eq. (S.30) the sign of $\delta D^{(\alpha)}$ is given by

$$\operatorname{sgn}\left[\delta D^{(\alpha)}\right] = -\operatorname{sgn}(\lambda)\operatorname{sgn}\left[\frac{\eta^{(\alpha)}}{k^{(\alpha)}} - \frac{\eta^{(\beta)}}{k^{(\beta)}}\right], \tag{S.34}$$

from which we can quantitatively define the terms 'target-aligned' and 'auxiliary-aligned': 'target-aligned' means $\eta^\alpha$ as a higher $(\alpha)$ content than $k^\alpha$ in the $(\alpha) - (\beta)$ plane, and $\left[\frac{\eta^{(\alpha)}}{k^{(\alpha)}} - \frac{\eta^{(\beta)}}{k^{(\beta)}}\right] > 0$; conversely, 'auxiliary-aligned' corresponds to $\left[\frac{\eta^{(\alpha)}}{k^{(\alpha)}} - \frac{\eta^{(\beta)}}{k^{(\beta)}}\right] < 0$. The change in $D^{(\alpha)}$ is a combination of the alignment and $\lambda$.

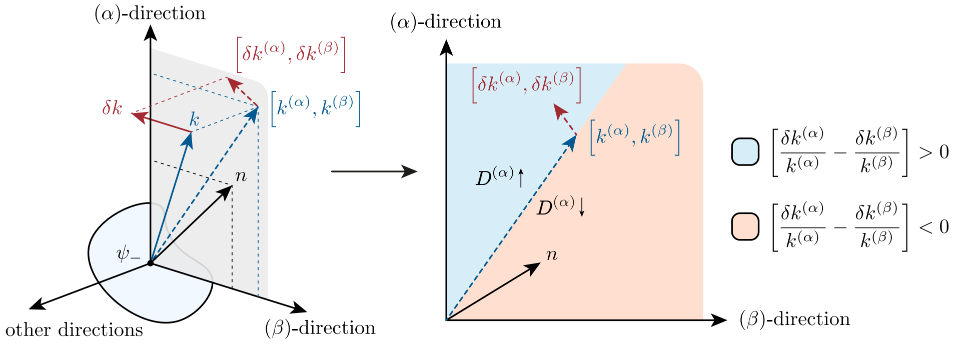


FIG. S2. **Relating changes in $D^{(\alpha)}$ to the phase space.** Under the assumption that $\partial_\alpha \partial_\beta f(\psi_-) = \partial_\alpha \partial_\beta f(\psi_+)$, the vector normal to the dilute phase boundary $n_\alpha$ stays constant, and changes in $D^{(\alpha)}$ depends solely on the spatial relationship between $\delta k^\alpha$, $n_\alpha$ and the axis $(\alpha)$. Left: the plane of interest (grey) is defined by the $(\alpha)$ axis and the $n_\alpha$ vector. The sign of $\delta D^{(\alpha)}$ then depends only on projections of $k^\alpha$ and $\delta k^\alpha$ onto this sub space, $[k^{(\alpha)}, k^{(\beta)}]$ (blue) and $[\delta k^{(\alpha)}, \delta k^{(\beta)}]$ (red).

#### V. EXPERIMENTAL PROCEDURE FOR G3BP1-RNA-SURAMIN PHASESCAN

The PhaseScan setup [2] is used for the experiment. G3BP1 protein with a green florescent protein (GFP) tag is produced using the same protocol as a previously published study [3], poly(A)-RNA is purchased from Sigma, and suramin is purchased from MedChemExpress. All three molecules are dissolved in the same buffer consisting of 100mM KCl, 2mM $MgCl_2$, 20mM PIPES, and 1% PEG (molecular weight 10 kilodaltons) at pH 7.2. To carry out the experiment, four aqueous inlets are used. The protein inlet contains 8uM G3BP1; the RNA inlet contains 200ng/µl RNA with 3µM Alexa546 florescent dye; and the suramin inlet contains 30µM suramin with 6µM Alexa647 florescent dye. The fourth inlet contains just buffer to maintain the total flow rates of all aqueous inlets to be 60µl/hr. The oil inlet contains HFE-7500 oil (FluoroChem) with 1% fluorosurfactant (RAN Bio), and a constant flow rate of 80µl/hr is used. 60,000 droplets are produced and quantified in the dataset.

The Alexa546 and Alexa647 dyes are added to quantify the total concentrations of RNA and suramin in each droplet using florescent channels corresponding to their wavelengths. For the protein, the 488nm wavelength channel is used. Total protein concentration in each droplet is calculated by taking the average intensity of all pixels in the droplet, and the dilute phase protein concentration is calculated by first computing the histogram of all pixel intensities in the droplet, and only using intensities that fall in the bottom 5-25% of the histogram to compute the average. We have also tried using the bottom 25-45% of the histogram but no significant difference is observed.

#### VI. FITTING OF LINE-SCAN DATA WITH TWO REGIMES

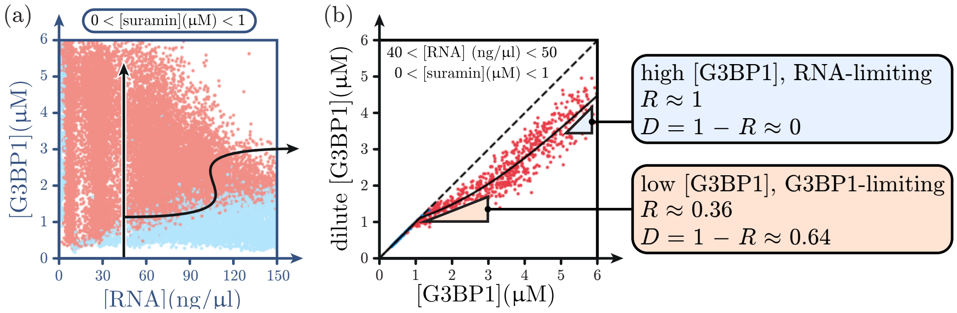


FIG. S3. **Two regimes of the homotypic line-scan.** (a) Illustration of the line-scan slice in the 2-D plane. (b) The homotypic response function has a gradient $R \approx 0.36$ at low [G3BP1] and $R \approx 1$ at high [G3BP1].

The 1-D line-scan data of G3BP1 has 3 different regions: 1 trivial homogeneous region, and 2 regimes in the phase separated region (Figure S3). At very low [G3BP1] the system is not phase-separated and a diagonal line is expected of the dilute versus total plot

$$y_1 = x. \tag{S.35}$$

As [G3BP1] increases, the onset of phase-separation results in a response that deviates from the diagonal and we locally expect a functional form

$$y_2 = ax + b. \tag{S.36}$$

Lastly at high [G3BP1] the response $R_1^1$ approaches 1, leading to

$$y_3 = x - c. \tag{S.37}$$

To introduce a smooth transitioning between $y_2$ and $y_3$ we construct the new function $y_{23}$ as

$$y_{23} = \frac{(y_1 + y_2) + \sqrt{(y_1 - y_2)^2 + \Delta^2}}{2} \tag{S.38}$$

where $\Delta$ serves as a smoothening parameter. To piece this with $y_1$, we simply use a parameter $x_0$ corresponding to the onset of phase-separation and write the final fitting function as

$$y_{\text{final}} = \begin{cases} y1 & \text{if } x < x_0 \\ y_{23} & \text{if } x \geq x_0 \end{cases} \tag{S.39}$$

and to ensure $y_{\text{final}}$ is continuous at $x_0$ we set the parameter $b$ such that $y_1(x_0) = y_{23}(x_0)$ given parameters $a$, $x_0$ and $c$. The response at $x_0$ is simply the derivative $y'_{\text{final}}(x_0)$.

With $x_0$ as a fitting parameter, it becomes possible to compute the phase boundary without directly imaging the sample and detecting the presence or absence of condensates. In the G3BP1-RNA-suramin PhaseScan we note that the phase boundary generated by detecting condensates in each droplet (Figure S4a to S4c) is very similar to that obtained from the critical [G3BP1] using the parameter fitting (Figure S4d). The condensate-dissolving effect of suramin and RNA is evident from the increasing trend of critical [G3BP1] with both suramin and RNA.

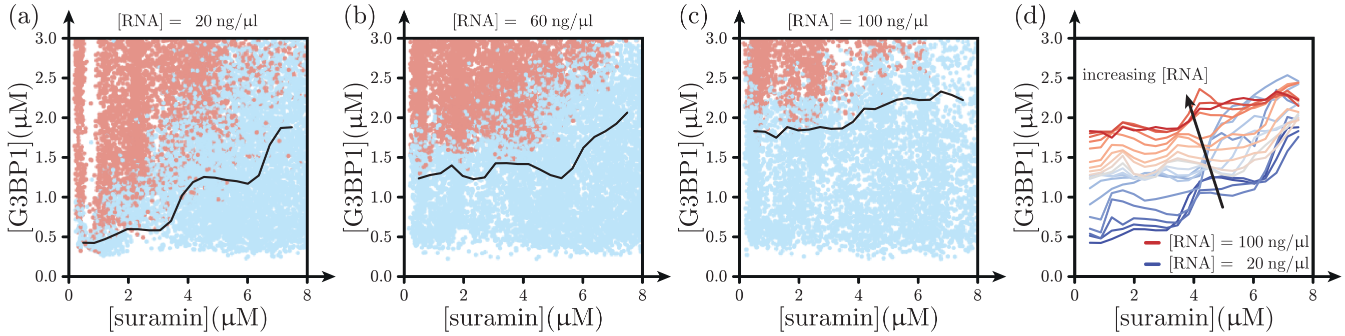


FIG. S4. **G3BP1 condensate dissolution effects of suramin and RNA** (a) to (c): at various [RNA] concentrations the G3BP1-suramin phase boundary shows an increasing trend. Scatter points are experimental data with image-based condensate detection, black solid line is the phase boundary estimated from fitting the line-scan response without condensate detection. (d) Estimated G3BP1-suramin phase boundary for a range of [RNA].

#### VII. DILUTE BAND BOUNDARY ANALYSIS

To provide further insight into the dilute phase band boundaries we illustrate how they arise from the multi-dimensional phase space. The 2-D G3BP1-RNA section probed using PhaseScan can be represented by a plane in the high-dimensional space (Figure S5a, top panel). The phase boundary exists as an $(N-1)$-dimensional surface, and it intersects the PhaseScan plane to produce a 1-D phase boundary (Figure S5a, middle panel). This is the ordinary phase boundary cross section seen in the experiment (Figure S5a, bottom panel). In effect, the set of data points populating the G3BP1-RNA section is sectioned by the phase boundary, and this sectioning idea is used in the following to visualise high-dimensional tie lines.

The dilute phase G3BP1 concentration measurement allows us to section the dataset further. We first pick an arbitrary concentration of G3BP1, for instance 3µM, and assign different shades of blue(red) to data points with(without) phase-separation depending on whether the dilute phase G3BP1 in that droplet is above or below 3µM. This plane of constant G3BP1 sections the high-dimensional phase space again. Outside the phase-separated region this sectioning is trivial since the dilute phase concentration is equivalent to the total concentration (Figure S5b, top panel) and the boundary observed in the 2-D PhaseScan plane is simply a horizontal line (Figure S5b, middle and bottom panels). To extend this sectioning into the phase-separated region, we note that the constant G3BP1 plane intersects the high-dimensional phase boundary and this intersection represents the set of points with dilute phase G3BP1 to be exactly 3µM. For any point within the phase separated region, its dilute phase composition resides on the phase boundary and the line connecting them is the tie line. As such, sectioning the phase separated region requires one to extrude the line of intersection between the constant G3BP1 plane and the phase boundary along tie lines (Figure S5c, top panel) and this procedure produces another $(N-1)$-dimensional surface. This newly created surface non-trivially sections the phase separated region and its intersection with the PhaseScan plane leads to the boundary between different shades of phase separated red points (Figure S5c, middle and bottom panels). Interestingly, points on this boundary satisfy the condition that their dilute phase G3BP1 is constant (3µM), and the boundary is by construction on the G3BP1-RNA plane, the gradient of the boundary is by definition $K$ [1]. Once the sectioning with a single G3BP1 concentration is established, this procedure can be easily performed for multiple threshold values and we represent the result as 'dilute phase bands' of alternating shades of blue and red (Figure S5d). These dilute phase bands are trivially horizontal outside the phase-separated region, and exhibit interesting shapes within. It is worth noting that although these bands arise from high-dimensional tie lines, they are curved in general even if the tie lines themselves are straight and parallel to one another. This is because the intersection between the constant G3BP1 plane and the phase boundary is in general curved, so the resulting sectioning surface is typically not flat.

Now we analyse the band boundaries, using indices 1 and 2 to denote G3BP1 and RNA respectively. In the low-RNA regime, the band boundaries appear to be straight and parallel to one another. We use a support vector machine (SVM) with a linear kernel to estimate the gradient of the band boundary at various levels of threshold [G3BP1] from 2 to 5 µM, and observe that $K$ stays constant within error, with $K = 0.047 \pm 0.005 \frac{\mu\text{M}}{\text{ng}/\mu\text{l}}$. Furthermore, we use the SVM to estimate $\frac{n_1}{n_2}$ in this regime. The result is $\frac{n_1}{n_2} = 0.17 \pm 0.02 \frac{\text{ng}/\mu\text{l}}{\mu\text{M}}$, and this can be combined with $K$ to give an estimate of $D^1$ for G3BP1. Noting [1]

$$D^1 = 1 - \frac{1}{1 + K\frac{n_1}{n_2}}. \tag{S.40}$$

we have $D^1 = 0.008 \pm 0.002$. The low $D$ value is apparent by observing that the phase boundary appears to be perpendicular to the RNA axis, so it is possible that RNA has a much larger $D$ in this regime. This can be understood as an RNA-limited situation, where the abundance of G3BP1 means each RNA molecule is bound by many G3BP1 that can cross-link between one another, and the G3BP1-covered RNA simply acts as a single-component phase separation system.

To relate $K$ to the G3BP1-RNA stoichiometry $\frac{k^1}{k^2}$, we use the relation

$$K = \frac{k^1}{k^2}\left[\frac{1}{1 + \sum_{(\alpha)=3}^{N} D^{(\alpha)}/D^{(2)}}\right] \tag{S.41}$$

and observe that $K$ will in general not be a constant across the phase space unless both $k^\alpha$ and $D^{(\alpha)}$ are constant. The constant $K$ seen in the experiment thus suggests a scenario where the RNA dominance $D^{(2)}$ is close to 1 and the tie line vectors are constant, allowing us to use the $K$ value as a stoichiometry indicator.

#### VIII. VALIDATION OF G3BP1-RNA CONDENSATE MODULATION BY SURAMIN

Microfluidic diffusional sizing of G3BP1 at various RNA and suramin concentrations is performed using the commercial Fluidity ONE-M machine. The basis of measurement is flowing two streams of solutions side by side under

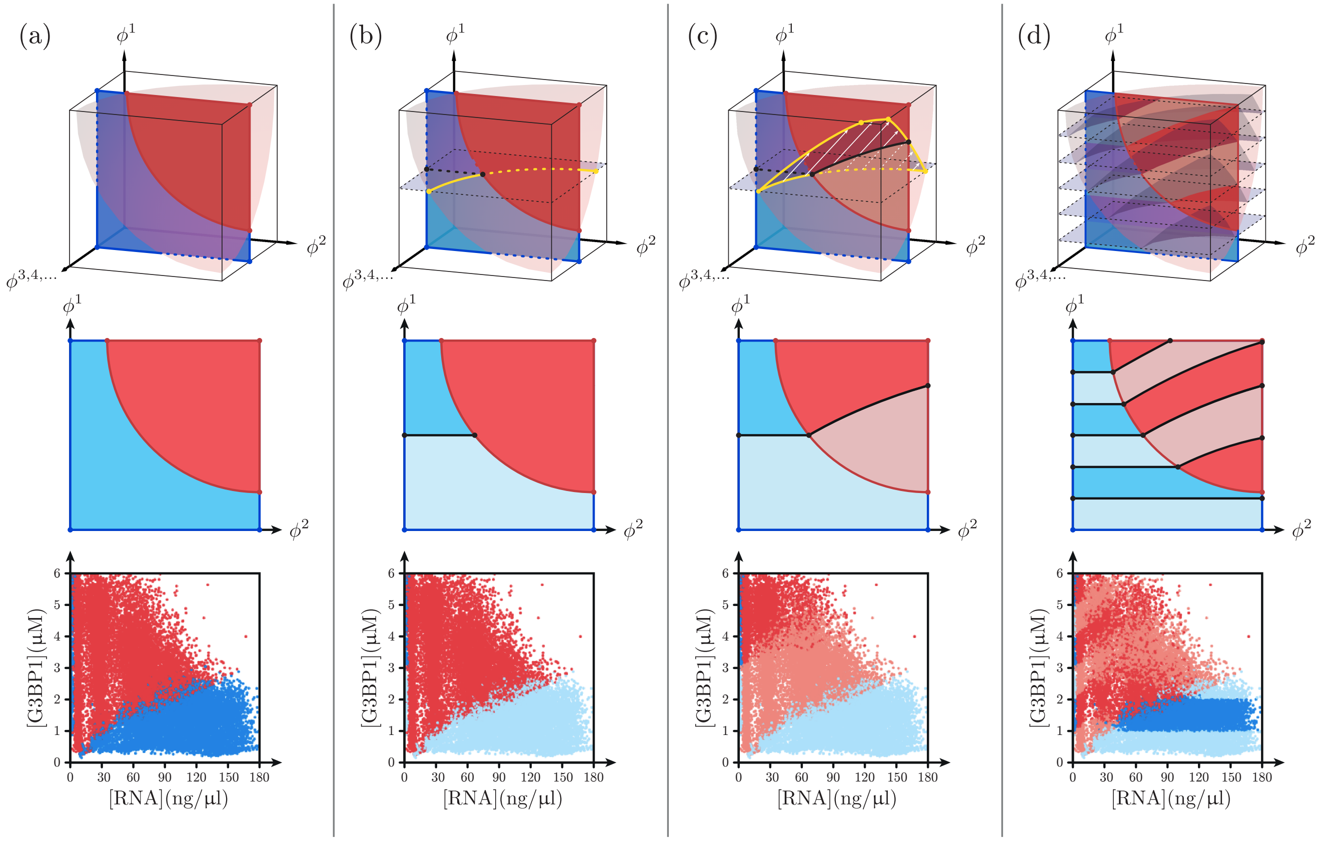


FIG. S5. **PhaseScan coupled to dilute phase concentration measurement efficiently probes the high-dimensional phase space.** (a) In the high-dimensional phase space (top panel) the phase boundary is represented as a transparent, curved surface, and a 2-D PhaseScan plane intersects the surface to produce a low-dimensional phase boundary observed in experiments (illustration in middle panel, data in bottom panel). (b) Sectioning the homogeneous space according to dilute phase concentration of solute 1 (G3BP1) leads to a horizontal boundary. Intersection between the sectioning plane and the phase boundary is highlighted with a yellow line in top panel. (c) This intersection extends into the phase separated region along tie line vectors (white arrows) to further section the high-dimensional space. This produces a dilute phase band boundary that is in general curved. (d) Multiple sectioning of the same space.

laminar conditions, with the protein present only in one of the streams. By measuring the concentration of protein that has diffused across the stream-stream interface the diffusion coefficient of G3BP1 can be determined [4].

The buffer solution contains 100mM KCl, 2mM $MgCl_2$, 20mM PIPES at pH 7.2. This is the same as that used in the PhaseScan experiment except no PEG is added to avoid phase separation. For each RNA and suramin condition the appropriate amount of RNA and suramin is added to both aqueous streams in the sizing machine while 1µM G3BP1 is added to only one stream so that the diffusion of G3BP1 can be quantified.

[1] D. Qian, H. Ausserwoger, T. Šneideris, R. Pappu, and T. P. J. Knowles, Dominance metric in multi-component binary phase equilibria, (2023).

[2] W. E. Arter, R. Qi, N. A. Erkamp, G. Krainer, K. Didi, T. J. Welsh, J. Acker, J. Nixon-Abell, S. Qamar, J. Guillén-Boixet, T. M. Franzmann, D. Kuster, A. A. Hyman, A. Borodavka, P. S. George-Hyslop, S. Alberti, and T. P. J. Knowles, Biomolecular condensate phase diagrams with a combinatorial microdroplet platform, Nature Communications **13**, 7845 (2022).

[3] D. Qian, T. J. Welsh, N. A. Erkamp, S. Qamar, J. Nixon-Abell, G. Krainer, P. St. George-Hyslop, T. C. T. Michaels, and T. P. J. Knowles, Tie-Line Analysis Reveals Interactions Driving Heteromolecular Condensate Formation, Physical Review X **12**, 041038 (2022), arXiv:2022.02.22.481401 [10.1101].

[4] J. P. Brody and P. Yager, Diffusion-based extraction in a microfabricated device, Sensors and Actuators, A: Physical **58**, 13 (1997).
